# Carrion flies as sentinels for monitoring rabbit calicivirus activity in Australia

**DOI:** 10.1101/585885

**Authors:** Robyn N Hall, Nina Huang, John Roberts, Tanja Strive

**Affiliations:** CSIRO Health & Biosecurity, Canberra, Australia; Centre for Invasive Species Solutions, Canberra, Australia; CSIRO Land and Water, Canberra, Australia

**Keywords:** calicivirus, RHDV, blowflies, surveillance

## Abstract

Rabbit caliciviruses are an essential tool for managing wild rabbit populations in Australia. Our understanding of rabbit calicivirus epidemiology in Australia currently depends on members of the public submitting liver samples from dead rabbits through a monitoring program called Rabbitscan. However, many wild rabbits die in inaccessible locations or are scavenged before sampling can occur, leading to considerable sampling bias. In this study we screened field-caught carrion flies for the presence of rabbit caliciviruses to monitor virus circulation patterns in the landscape, with an aim to establish a less biased epidemiological surveillance tool. Carrion flies were collected from two study sites over a 22 month period and these samples were used to optimise and validate molecular testing methods in this sample type for the currently circulating rabbit calicivirus variants. Virus was clearly detectable in field-caught carrion flies using optimised SYBR-green RT-qPCR and RT-PCR assays. However, variant identification was frequently hindered by the low virus loads present in carrion fly samples and spurious RT-PCR amplification. This was overcome by frequent sampling, which effectively acts as replicate sampling to verify inconclusive results. There was good correlation between virus detections in carrion flies and in samples recovered from wild rabbits, both temporally and for virus variant identification. The methods reported here provide a robust and efficient additional surveillance tool to monitor rabbit calicivirus activity at a landscape scale, which in turn can help to guide more effective rabbit management programs.

## Introduction

Wild European rabbits (*Oryctolagus cuniculus*) are one of Australia’s most invasive vertebrate pests, directly threatening over 300 native plant and animal species and costing Australian agriculture over $200 million AUD annually. Wild rabbits are currently controlled using an integrated approach involving both conventional pest management methods, such as shooting and poisoning, and biocontrol agents, principally rabbit caliciviruses (Le Pendu et al., 2017). It is feasible that synergism between virus variants could be exploited to maximise the effectiveness of biocontrol programs, however this requires knowledge of prior infection history in different wild rabbit populations. Five pathogenic rabbit caliciviruses have thus far been reported in Australia: 1) the original V-351 RHDV (GI.1c) deliberately released as a biocontrol agent in the mid-1990s, and its descendants; 2) an additional rabbit biocontrol agent and released nationwide in March 2017 (Invasive Animals CRC, 2014); 3) a GI.1a variant, later identified as a recombinant GI.4e-GI.1a (subsequently referred to as GI.1a-Aus) (Mahar et al., 2018); 4) a recombinant GI.1b-GI.2 virus (also known as RHDV2 or RHDVb) (Hall et al., 2015); and 5) a recombinant GI.4e-GI.2 virus, comprising the non-structural genes of GI.1a-Aus and the structural genes of GI.2 (Hall et al., 2018).

Both molecular and serological methods have previously been used to monitor rabbit calicivirus activity across Australia. Molecular methods are relatively quick, accurate, and sensitive, however, sample collection is extremely biased, with most wild rabbits dying in inaccessible locations and/or being removed by scavengers before collection. Serological testing works well as a population tool to identify previous exposure of populations to rabbit caliciviruses (Cooke et al., 2000; Cox et al., 2017). However, the confidence and power of the conclusions drawn from results of serological testing depends strongly on the sample size, and collection of shot samples from multiple animals requires considerable time and person resources. Furthermore, serological tools are only able to broadly distinguish between GI.1 and GI.2 viruses, and accurate identification to the variant level and of mixed infections is challenging due to cross-reactivity between assays (Strive et al, in preparation). Due to the limitations of existing methods, a less biased and more systematic sampling method is required for surveillance of rabbit calicivirus activity across Australia, firstly to determine the spatial and temporal distribution of specific virus variants and subsequently to infer the interactions between the different viruses.

Carrion flies have long been known to be able to mechanically transmit rabbit caliciviruses (Asgari et al., 1998; Cooke and Fenner, 2002; McColl et al., 2002; Henning et al., 2005; Schwensow et al., 2014). Laboratory studies have detected rabbit calicivirus (GI.1c) by RT-PCR in various fly species (*Calliphora, Chrysomya, Hydrotaea, Lucilia, Musca, Sarcophaga*, and *Oxysarcodextia* species), as well as in two species of *Aedes* mosquitos (Asgari et al., 1998; McColl et al., 2002; Henning et al., 2005). It was further shown that virus was detectable in flies for more than 11 days, however, virus was only detectable on the legs of flies for 7 hours (Asgari et al., 1998). Virus has also been detected in flyspots (faecal and regurgitation spots) by RT-PCR, and it has been demonstrated that flyspots contained sufficient virus to cause disease when used to infect susceptible rabbits by oral infection (Asgari et al., 1998). Transmission trials have also shown that bush flies (*Musca vetustissima*) fed on carcasses of GI.1c-infected rabbits were able to naturally transmit infection to susceptible rabbits (McColl et al., 2002).

Since carrion flies are relatively easy and inexpensive to collect, they could prove a valuable additional systematic sampling tool for landscape scale monitoring of RHDV activity using high-sensitivity molecular detection methods. As the genetic diversity of rabbit caliciviruses has greatly increased in Australia in recent years, and several virus variants may co-circulate at the same time, the aim of this study was to develop molecular tools and protocols to reliably detect circulating rabbit caliciviruses in carrion flies to the virus variant level. These tools were subsequently validated by analysing carrion flies sampled at two sites closely monitored for rabbit calicivirus activity over a period of 22 months.

## Materials and methods

### Fly collection

Carrion flies were collected approximately weekly between 26 September 2016 and 30 June 2018 at two locations, Murrumbateman NSW (−35.00, 149.02) and Black Mountain ACT (−35.27, 149.11). Flies were trapped initially in LuciLure traps (BioGlobal, Australia) with commercial attractant and later, for convenience, in Envirosafe™ fly traps (Bunnings, Australia) baited with either Envirosafe™ fly attractant (Bunnings, Australia), chicken liver, or commercial attractant (BioGlobal, Australia), or a combination of the above. Attractant was placed in specimen jars covered with a gauze swab to prevent flies coming into direct contact with the bait (Figure 1). Traps were left out for one to seven days and upon collection were briefly frozen at −20 °C to immobilise flies before flies were transferred to storage containers at −20 °C. Fly traps and all components (except bait) were soaked in 10% household bleach for 30 minutes before being rinsed and re-used.

**Figure 1:**
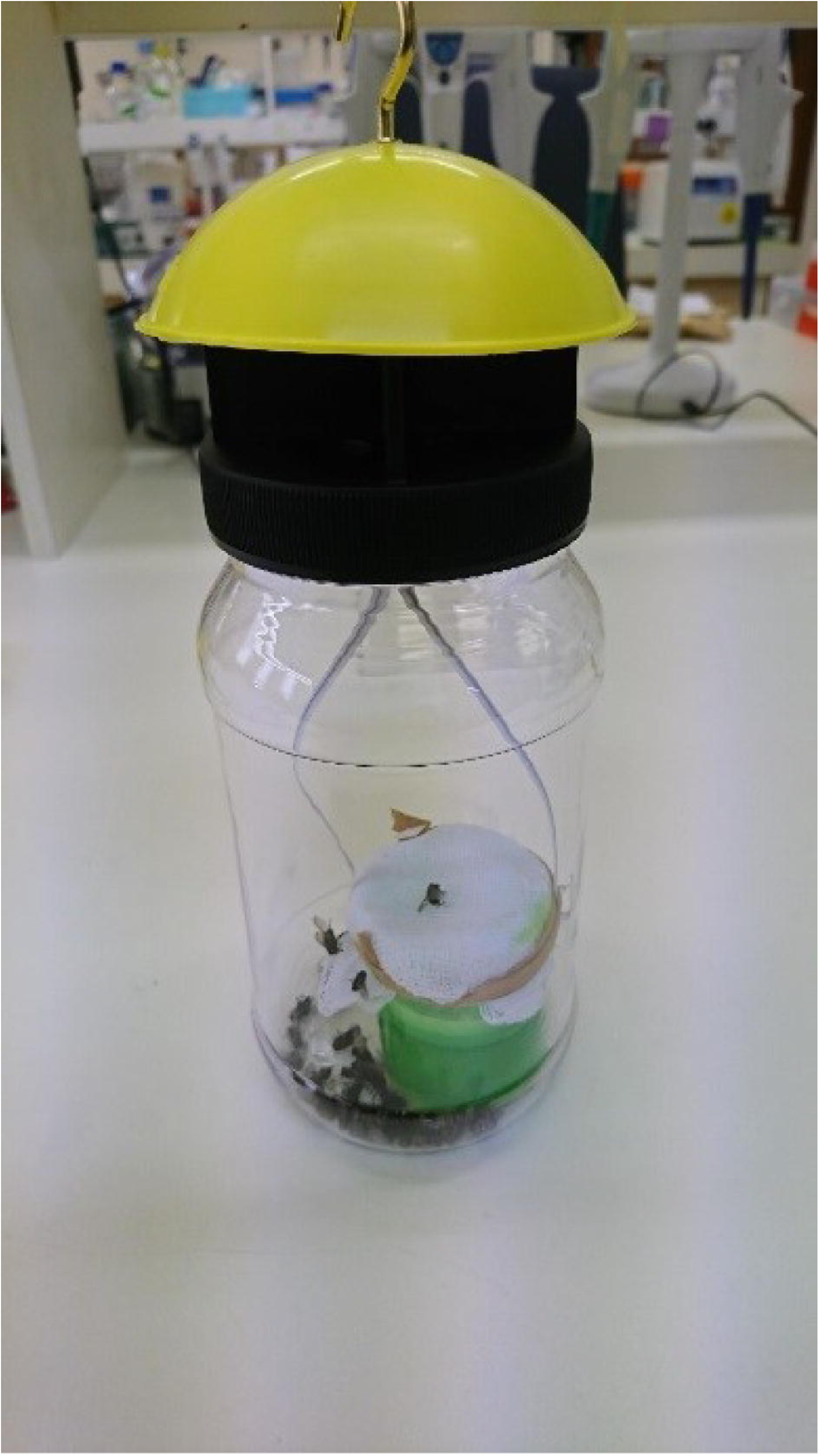
Envirosafe™ fly trap containing commercial bait. To prevent flies contacting the bait the specimen jar was covered by a gauze swab.

### Optimisation of pooling strategy

To optimise the pooling strategy (number and species of flies per pool) an early field collection (Black Mountain 25/11/2016) that contained a large number of flies and was positive for GI.4e-GI.2 rabbit calicivirus was used. We elected not to use laboratory-spiked samples since this field collection more accurately represents the characteristics of our final target collections.

To determine the effect of fly species on viral load and analytical sensitivity, four to five flies of five different species (based on morphological identification) were examined individually for the presence of virus. RNA was extracted and RT-qPCR and RT-PCR were conducted as described below. Species was confirmed by amplifying a region of mitochondrial DNA using the primers C1-J-2495 and C1-N-2800 (Wells and Sperling, 2001) and Platinum Taq DNA polymerase (Life Technologies) as per manufacturer’s instructions. Sanger sequencing was then conducted at the Australian Cancer Research Foundation Biomolecular Resource Facility, and sequences were analysed for the closest genetic match using NCBI Blastn against the nr database as implemented in Geneious 11.1.

To determine the effect of fly number on analytical sensitivity, we prepared five replicate pools containing either one, three, five, or ten flies (mixed species) per pool. RNA was extracted and RT-qPCR and RT-PCR were conducted as described below.

### Optimisation of RNA extraction method

To determine the effect of alternative homogenisation buffers and pre-extraction freeze-drying on analytical sensitivity, we prepared 12 replicate pools containing ten flies (mixed species) per pool. RNA was extracted from pools as follows: three pools using the standard protocol described below; three pools using a modified protocol incorporating freeze-drying of flies; three pools using the standard protocol with an alternative 10 mM Tris-HCl pH 8.5 homogenisation buffer containing 10 μl.ml^-1^ β-mercaptoethanol; and three pools using the standard protocol with an alternative phosphate buffered saline (PBS) homogenisation buffer containing 10 μl.ml^-1^ β-mercaptoethanol.

RNA was extracted using the Maxwell RSC simplyRNA tissue kit (Promega, Alexandria, NSW) on a Maxwell RSC system (Promega). Briefly, to each sample 10 volumes (minimum 200 μl) of homogenisation buffer was added per mg of fly weight. Samples were heated at 70 °C for 2 minutes before homogenisation using a Precellys 24-dual tissue homogeniser (Bertin Technologies, Montigny-le-Bretonneux, France). Samples were clarified at 3,000 *g* for 3 minutes and homogenate (minimum 200 μl) was combined with an equal volume of lysis buffer and mixed by vortexing. This homogenate/lysis buffer mix (400 μl) was used for extraction. The complete protocol is available at dx.doi.org/10.17504/protocols.io.ux7exrn. Known rabbit calicivirus negative flies (*Calliphora augur* reared in clean conditions in the laboratory) were extracted in parallel with each set of RNA extractions as a negative extraction control.

This protocol was modified as follows when flies were freeze-dried. Samples were freeze-dried overnight in a Flexi-Dry MP freeze-dryer (FTS Systems, Stone Ridge, New York). Once dry, samples were homogenised using a Precellys 24-dual tissue homogeniser (Bertin Technologies). Ten volumes of homogenisation buffer was then added per mg of dry fly tissue. Samples were heated, clarified, and mixed with lysis buffer as described above.

### All RNAs were stored at −80 °C

#### RT-qPCR and endpoint strain-specific PCRs

To compare analytical sensitivity during validation, viral loads were quantified using a SYBR green based RT qPCR for the generic detection of all Australian rabbit caliciviruses, as described previously (Hall et al., 2018). Detections were confirmed using the strain-specific lagovirus multiplex RT-PCR assay described previously (Hall et al., 2018). RNAs were diluted 1/10 in nuclease-free water prior to RT-PCR and were used undiluted as template in the RT-qPCR assay.

For routine monitoring of field-caught fly samples, a serial testing procedure was used where samples were first run through the universal RT-qPCR assay and those samples with virus loads greater than 1000 capsid gene copies per μl of RNA (cut-off derived during validation) were subsequently analysed using the strain-specific RT-PCR assays in singleplex format for virus variant identification.

#### Strain identification in rabbits and hares

Liver samples from rabbits and hares found dead at Black Mountain or Murrumbateman were screened for rabbit caliciviruses using previously reported methods (Hall et al., 2018). No animal ethics permit is required in Australia for sample collection from rabbits or hares that are found dead.

## Results

A series of experiments was conducted to validate and optimise molecular testing protocols for the strain-specific detection of rabbit caliciviruses in carrion flies. Specifically, we focussed on optimising the pooling strategy (number and species of flies from individual traps) and RNA extraction method using flies from a single known-positive field collection. We then assessed the suitability of existing RT-qPCR and RT-PCR assays for detection of rabbit caliciviruses in fly RNA. These existing assays were developed for use on liver RNA, although other sample types including bone marrow, whole blood, maggots, and skin have also been used for testing and returned positive results (unpublished data). Once an optimal test procedure was determined, we used these protocols to monitor rabbit calicivirus circulation in the environment using carrion flies over a 22 month period at two locations, Murrumbateman (NSW) and Black Mountain (ACT), approximately 35 km apart.

### Species or number of flies does not affect sensitivity of detection

Five different fly species were tested to determine whether virus load correlated with species. Individuals were identified morphologically as either *Calliphora augur, C. canimicans bezzii, C. stygia, Chrysomya rufifacies*, or *Chrysomya varipes* (Wallman, 2001). A region of the mitochondrial DNA of these individuals was also sequenced to verify species identity. Species was confirmed in most cases, with a few exceptions. One individual morphologically identified as *C. stygia* was genetically classified as *C. hilli hilli* and all *C. canimicans bezzii* were genetically identified as *C. ochracea*. Morphologically, *C. ochracea* is quite distinctive and further investigation revealed that no sequences for *C. canimicans bezzii* are present in GenBank. Therefore, this classification of *C. ochracea* is likely to be incorrect. These individuals were subsequently referred to as *Calliphora* (species unidentified).

Every individual fly apart from one *Chrysomya rufifacies* was positive on both RT-qPCR and strain-specific RT-PCR (Figure 2a). There was no significant difference in the viral load carried by different fly species. To test whether the number of flies used per RNA extraction affected the sensitivity of detection five replicate pools of either three, five, or ten individual flies of mixed species were examined by RT-qPCR. Again, no significant differences in viral loads were observed in the different pool sizes, and the viral loads in pooled samples were comparable to those detected in individual flies (Figure 2b).

**Figure 2:**
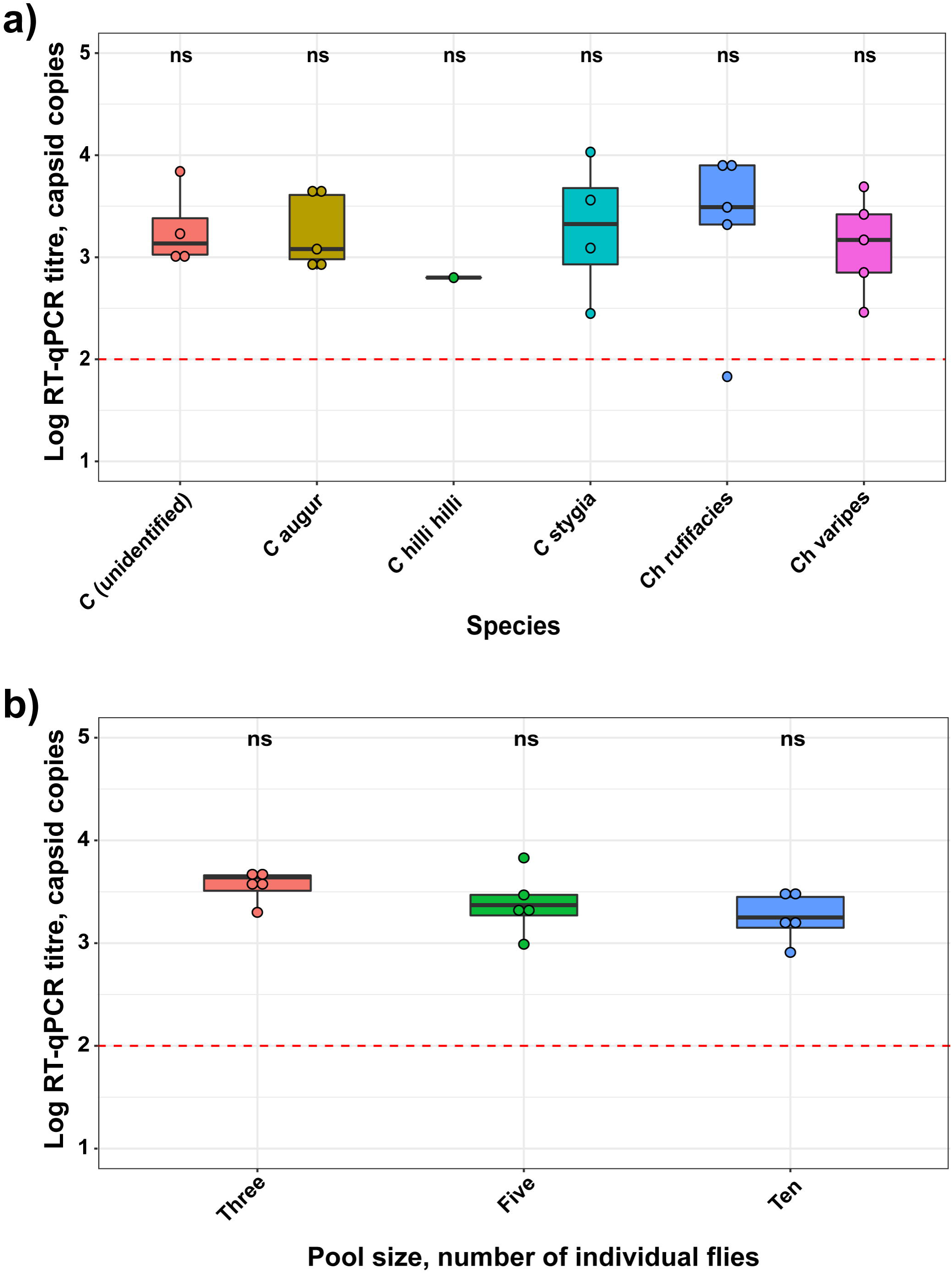
Four to five individual flies of each species (a) or five replicate pools of *n* = 3, 5, or 10 flies (b) were tested for the presence of rabbit caliciviruses using RT-qPCR. Statistical comparisons between species were performed using pairwise Wilcoxon tests in the R package “ggpubr” as implemented in R3.5.1. The red dashed line indicates the limit of quantification of the RT-qPCR assay.

### Optimisation of RNA extraction protocols for rabbit calicivirus detection in carrion flies

When processing larger pools of flies it became apparent that for increased throughput an alternative homogenisation buffer would be required, since the volume of commercial homogenisation buffer provided in the kit was insufficient. We therefore compared two alternative homogenisation solutions, either 10 mM Tris-HCl pH 8.5 or 1× PBS, both containing 10 μl.ml^-1^ β-mercaptoethanol to inactivate environmental RNases, with the commercial buffer. We also assessed the suitability of freeze-drying flies prior to RNA extraction, in order to quantitate virus load on a dry matter basis. No significant differences in viral load were observed between freeze-dried or fresh flies or between the alternative homogenisation buffers used (Figure 3).

**Figure 3:**
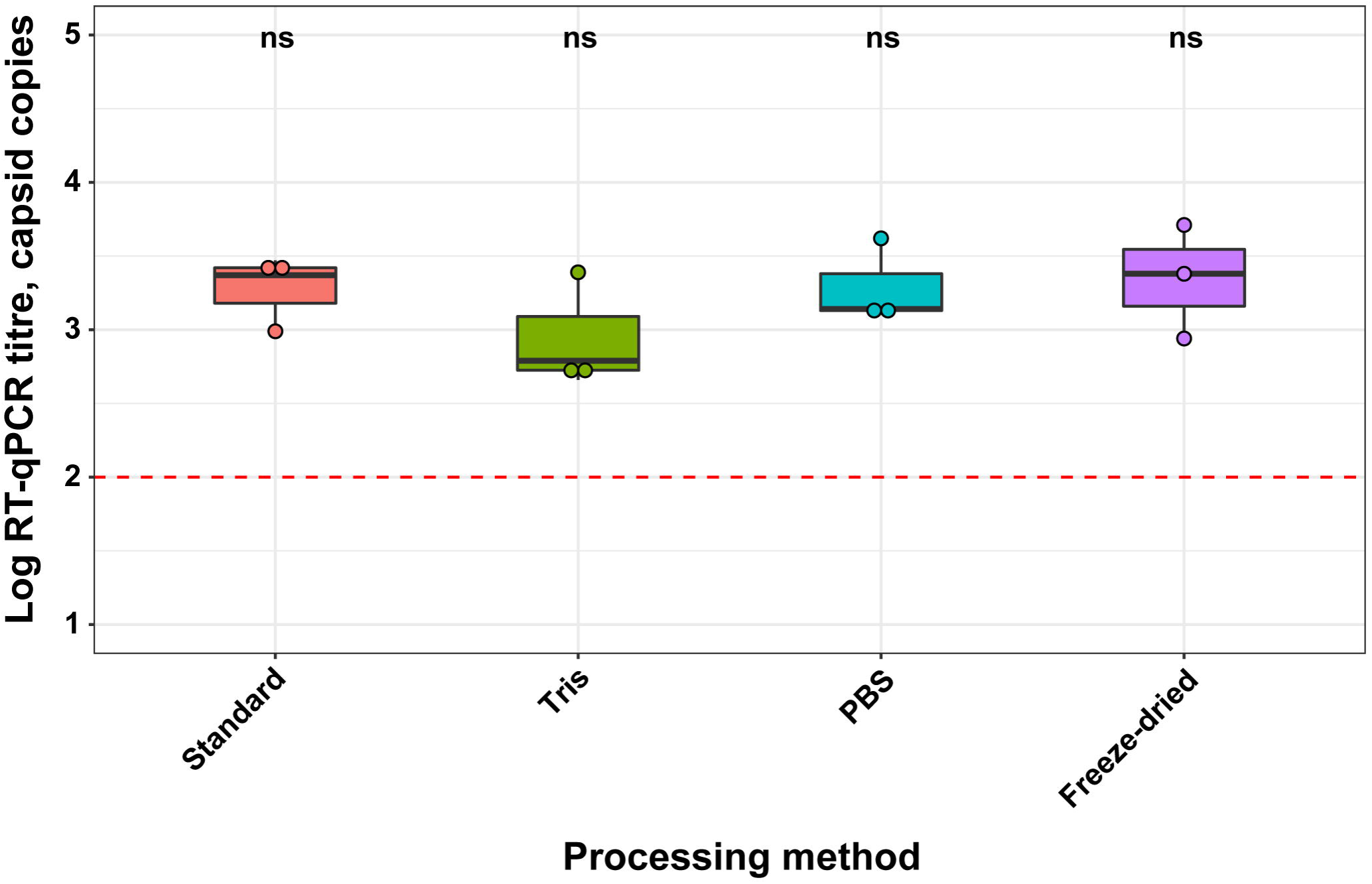
Total RNA was extracted from three replicate pools of ten flies by either the standard extraction method, substituting the homogenisation buffer for 10mM Tris + 10 μl.ml^-^ 1 β-mercaptoethanol or 1x PBS + 10 μl.ml^-1^ β-mercaptoethanol, or after freeze-drying pools overnight. Virus load was quantified using RT-qPCR. Statistical comparisons between species were performed using pairwise Wilcoxon tests in the R package “ggpubr” as implemented in R3.5.1. The red dashed line indicates the limit of quantification of the RT-qPCR assay.

### Modification of existing RT-qPCR and RT-PCR assays enabled robust detection of rabbit caliciviruses in carrion flies to the virus variant level

An optimised RNA extraction protocol that included freeze-drying of flies was then used to process 99 fly pools collected between September 2016 and June 2018 at two locations near Canberra ACT. These were screened in series as previously recommended (R. N. Hall et al., 2018)—initially samples were tested in a universal lagovirus SYBR-green-based RT-qPCR and those above the limit of quantification of the assay (i.e. 100 capsid copies per μl of RNA) were subtyped using a strain-specific RT-PCR. Of the 99 pools collected over the sampling period, 49 (49%) samples had viral loads above this limit of quantification. When using the RT-PCR in multiplex format, frequently non-specific bands were present that made interpretation very challenging, despite diluting the RNA template to mitigate the effects of PCR inhibitors that may be present. To try to avoid the formation of these non-specific bands, subtyping was performed using singleplex RT-PCR assays for each virus variant (i.e. GI.1, GI.2, GI.1a-K5, GI.1a-Aus). Based on these singleplex endpoint RT-PCR results, a virus strain was unambiguously assigned for 24 of these 49 samples (49%) (Figure 4). The remaining 25 samples continued to return inconclusive results, either returning negative results on RT-PCR despite being positive on RT-qPCR, inconsistent results on repeated testing, or only amplifying very weak bands. Contrastingly, on rabbit liver samples strain determination is extremely accurate on samples with viral loads greater than 100 capsid copies per μl of RNA (R. N. Hall et al., 2018). Increasing the RT-qPCR cut-off to 1000 capsid copies per μl of RNA increased the stringency of testing, with 26 samples testing positive. Of these samples, strain was unambiguously determined in 18 (69%) of those samples.

**Figure 4:**
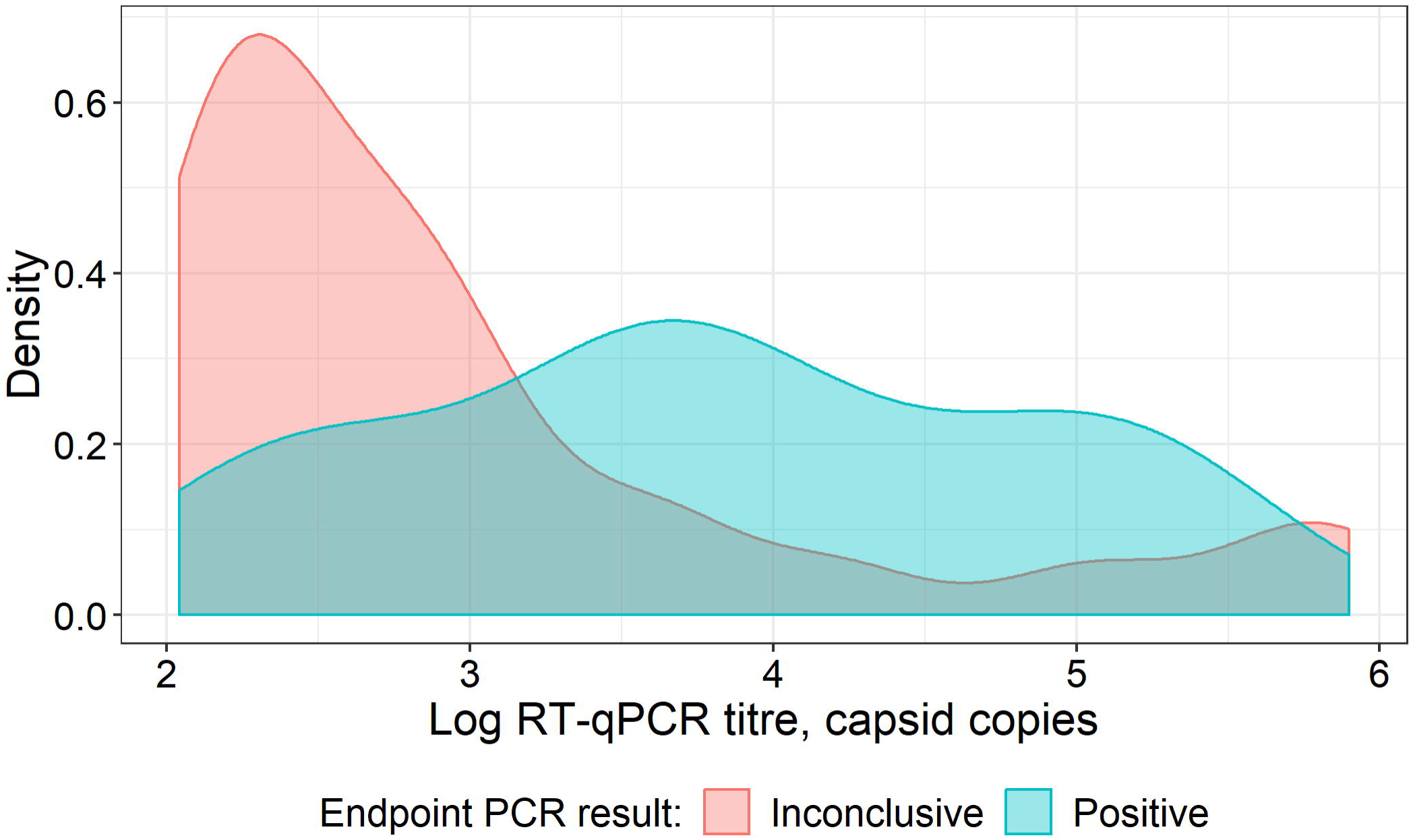
Density plots were used to compare the distributions of positive and inconclusive fly samples as a function of viral load as determined by RT-qPCR. Flies were collected between September 2016 and June 2018 and were screened by a universal lagovirus SYBR-green-based RT-qPCR. Those above the limit of quantification of the assay (i.e. 100 capsid copies per μl of RNA) were subtyped using strain-specific endpoint RT-PCR assays. Samples for which strain was unambiguously assigned were classified as positive while those that were negative on endpoint RT-PCR, gave inconsistent results on repeated testing, or only amplified very weak bands were classified as inconclusive.

### Carrion flies provide a cheap and efficient method to monitor rabbit calicivirus outbreaks

Using the modified RT-qPCR threshold value of 1000 capsid copies per μl of RNA, viral loads were plotted as a time series to determine whether carrion flies were suitable proxies for detecting rabbit calicivirus circulation in the environment. Distinct calicivirus-positive fly collections were observed over the sampling period based on this fly sampling method (Figure 5). Positive samples were observed at Black Mountain in December 2016, April 2017, October 2017, December 2017-January 2018, and April 2018. Similarly, positive fly samples were observed at Murrumbateman in April 2017, October 2017, January-February 2018, and March-April 2018. Murrumbateman potentially also had a positive sample in December 2016, however, the virus variant could not be determined conclusively. At both locations, calicivirus circulation detected via fly sampling correlated closely with detection of calicivirus-positive dead lagomorphs (i.e. rabbits and hares), both temporally and for virus variant identification (Figure 5). At Black Mountain calicivirus-positive lagomorphs were detected in April-May 2017, October 2017, January 2018, and May-June 2018, while at Murrumbateman they were detected in April-June 2017, October-November 2017, and June 2018.

**Figure 5:**
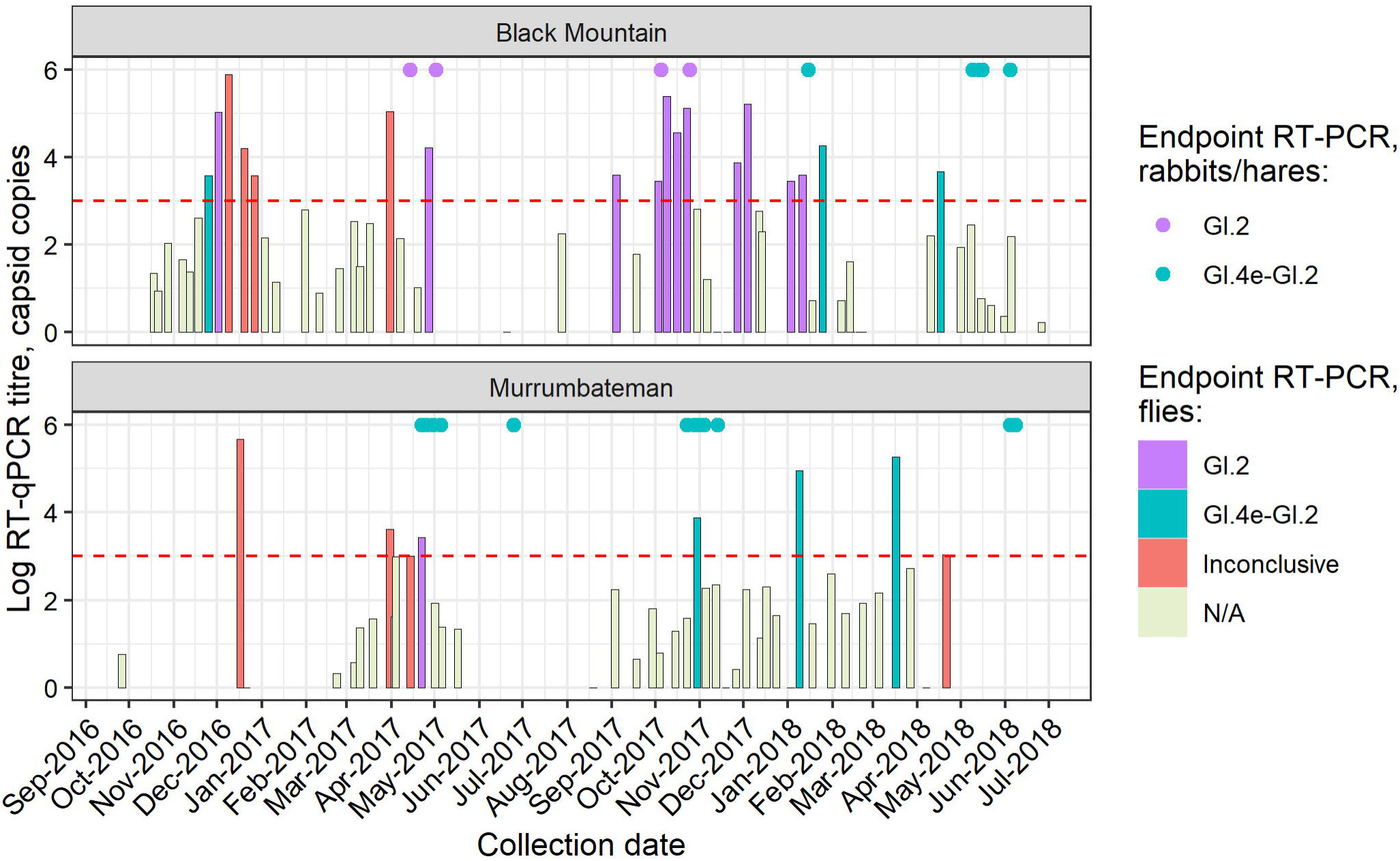
Flies were collected regularly from two locations around the ACT between September 2016 and June 2018 to monitor rabbit calicivirus circulation in the environment. Total RNA was extracted and viral load was quantified by RT-qPCR. For samples with viral loads >1000 capsid copies per μl of RNA, virus variant was determined by strain-specific endpoint RT-PCR. Lagomorphs (i.e. rabbits or hares) found dead at these locations over the sampling period and that were positive for rabbit calicivirus were plotted as points at 1 × 10^6^ capsid copies per μl of RNA for visualisation purposes.

## Discussion

Monitoring rabbit calicivirus circulation across Australia is important both to inform management of wild rabbit populations and control and prevention of calicivirus outbreaks in domestic rabbits. Knowing the geographical and temporal distribution of different caliciviruses can help to 1) guide the development, and 2) increase the effectiveness of improved rabbit management programs, while domestic rabbit owners can instigate targeted control and prevention measures when calicivirus is active in their geographical region. We have demonstrated a novel sampling strategy to enable robust and efficient identification of caliciviruses to the virus variant level circulating in Australian lagomorph populations.

There was no significant difference in detection sensitivity between the fly species tested, namely *Calliphora augur, C. hilli hilli, C. stygia, Chrysomya rufifacies, Chrysomya varipes* or an unidentified calliphorid species (presumably *C. canimicans bezzii* based on morphological identification). All individuals were obtained from a single collection, and viral RNA was detected in amounts greater than the limit of detection of the assay in all flies tested except a single *Chrysomya rufifacies*. Since it is unlikely that 100% of flies carry rabbit calicivirus at any given point in time, this suggests that contamination between flies occurs within a trap, probably through contact with flyspots excreted from infected flies (Asgari et al., 1998). Detection sensitivity was also found to be independent of the number of individual flies pooled for RNA extraction, at least for pool sizes between one and ten individuals. Presumably, this is due to the virus being concentrated in the digestive tract of flies combined with a dilution effect when additional fly tissue is added. Our choice of pool size now depends on the minimum tissue amount recommended for RNA extraction (20 mg dry fly tissue) and a maximum practical amount for processing (approximately 100 mg of dry fly tissue). Additionally, no significant differences in detection sensitivity were observed when flies were freeze-dried prior to processing, or when alternative buffers were used for tissue homogenisation. Freeze-drying flies facilitates quantification of viral load in flies on a dry matter basis, since flies can vary considerably in their moisture content depending on environmental conditions. Homogenisation is also more thorough for freeze-dried samples, compared to fresh frozen samples. Flies stored in saturated salt solutions (e.g. RNALater) should not be freeze-dried, as the salt crystallises and interferes with quantification and homogenisation.

We then used previously described RT-qPCR and strain-specific endpoint RT-PCR assays to monitor rabbit calicivirus circulation in carrion flies over a 22 month period at two locations approximately 35 km apart near Canberra ACT. By testing carrion fly samples in series, initially with the RT-qPCR assay and subsequently with the strain-specific RT-PCR assays for samples >1000 capsid copies per μl of RNA, we were able to identify clear temporal peaks of rabbit calicivirus activity at both sampling locations. Virus variant identification was hindered by a relatively low positive predictive value of the SYBR-green-based RT-qPCR assay used here, and more sensitive and specific tests may be able to further improve this testing pipeline in the future. Depending on the relative importance of false negative versus false positive results, this threshold for strain identification can be adjusted as required. Testing of multiple replicate pools from a single trap or multiple collections from a specific time point and location may also increase detection sensitivity and specificity, however, this must be balanced against the costs incurred with additional testing.

The pattern of positive fly samples was very similar at both Black Mountain and Murrumbateman, with peaks of virus detection in March-April, October, and December-January. This similarity is not surprising, given that the distance between these locations is only 35 km and rabbit caliciviruses can spread at rates of up to 400 km per month during autumn and spring (Kovaliski, 1998). Furthermore, flies are known to move up to 15 km per day (cited in McColl et al., 2002). There was good correlation between peaks of detections in flies and detections in rabbits, at the temporal, geographic, and virus variant levels. Virus was detected in both rabbits and flies in April 2017, October 2017, and January 2018 at Black Mountain and in April and October 2017 at Murrumbateman. Detections in carrion flies typically preceded detection in lagomorphs by around two weeks. The virus detected in flies matched that detected in lagomorphs except during April 2017 at Murrumbateman, where recombinant GI.4e-GI.2 was detected in lagomorph carcasses while GI.2 was detected in flies. All subsequent detections from Murrumbateman were recombinant GI.4e-GI.2.

Virus was detected in flies only at Black Mountain in December 2016 and April 2018 and Murrumbateman in January and April 2018. This suggests that using carrion flies as proxies to monitor rabbit calicivirus circulation in the environment may in fact be more sensitive than relying on detection of rabbit carcasses. Interestingly, of the two fly samples collected from Black Mountain in December 2016, one was identified as GI.4e-GI.2 while the other was identified as GI.2. Previous detections of calicivirus in rabbit carcasses at Black Mountain were all GI.2, however, recombinant GI.4e-GI.2 was detected at one location in the ACT in a dead rabbit approximately 13 km away in December 2016. It is possible that this detection at Black Mountain was in fact dispersal of flies from this distant site, or that the two viruses were co-circulating and no rabbit carcass with the recombinant GI.4e-GI.2 was detected. Virus was detected in rabbits only in May 2018 at Black Mountain, which may have been related to the April 2018 detections in flies. Additionally, virus was detected in rabbits only in June 2017 and 2018 in Murrumbateman. Fly samples were challenging to obtain during the winter months, when fly activity is low in this region of Australia. Indeed, no flies were trapped from Murrumbateman during June-July 2017 or May-June 2018. Consideration should be given to an alternative sample source when flies are not active in a given region, for example, over winter.

It should be noted that not all fly samples collected during outbreaks were positive. There is a high rate of negative or inconclusive samples, particularly when using the stringent threshold we have selected. This hampers the interpretation of single fly collections. This suggests that sampling of carrion flies to monitor calicivirus activity may be best suited for temporal sampling, with samples collected regularly, and probably not less than once a month, although sampling frequency for optimal monitoring has not been modelled or experimentally determined at this stage. To function as a tool to guide wild management control programs and disease control and prevention programs in domestic rabbits, regular long-term sampling would be required. To instigate a national systematic sampling network, the spatial scales at which calicivirus dynamics change in flies still needs to be investigated. The fly sampling method reported here provides a robust and efficient additional method to monitor broad-scale epidemiological patterns of rabbit calicivirus activity that could be used to help guide wild rabbit management programs.

## Acknowledgements

The authors wish to thank Roslyn Mourant, Lily Tran, and Gordon Soon for their assistance with sample collection and processing and Adam Croxford for helpful suggestions. We also thank Peter Kerr and Amanda Padovan for critical review of the draft manuscript. This project was funded by the Commonwealth Scientific and Industrial Research Organisation— Health and Biosecurity.

